# Tetanizing wakeful consolidation: ten-hertz repetitive visual stimulation enhances the offline gain of visual learning

**DOI:** 10.1101/2024.09.20.614122

**Authors:** Xin-Yue Yang, Chuyue Zhao, Zhentao Zuo, Aming Li, Huan Luo, Qing He, Fang Fang

**Affiliations:** School of Psychological and Cognitive Sciences and Beijing Key Laboratory of Behavior and Mental Health, Peking University, Beijing, China; Key Laboratory of Machine Perception (Ministry of Education), Peking University, Beijing, China; Peking-Tsinghua Center for Life Sciences, Peking University, Beijing, China; IDG/McGovern Institute for Brain Research, Peking University, Beijing, China; State Key Laboratory of Brain and Cognitive Science, Institute of Biophysics, Chinese Academy of Sciences, Beijing, China; University of Chinese Academy of Sciences, Beijing, China; Center for Systems and Control, College of Engineering, Peking University, Beijing, China; Center for Multi-Agent Research, Institute for Artificial Intelligence, Peking University, Beijing, China

## Abstract

Consolidation of encoded information is vital for learning and memory, often explored during sleep. However, the consolidation during post-encoding offline wakefulness remains largely uncharted, especially regarding its modulation and brain mechanisms. Here, we unraveled frequency-dependent modulatory effects of repetitive visual stimulation (RVS) on wakeful consolidation of visual learning and investigated the underlying neural substrates. After training on an orientation discrimination task, exposure to 10-Hz grating-form RVS enhanced, while 1-Hz RVS deteriorated, the discrimination performance in a subsequent retest. However, 10-Hz uniform-disk RVS failed to facilitate wakeful consolidation, suggesting that alpha entrainment alone was not the facilitative mechanism. Using neuroimaging of multiple modalities, we observed augmented event-related potential and heightened neural excitation in the early visual cortex after 10-Hz grating-form RVS, implying an involvement of long-term potentiation-like (LTP-like) plasticity. Collectively, we provide a new photic method for modulating the offline processing of encoded sensory information and suggest a role of sensory tetanization in the modulation.

## Introduction

Learning and memory encompass encoding, consolidation, and retrieval processes that necessitate neural plasticity^1,2^. Neural plasticity associated with learning and memory has been predominantly studied during encoding. For example in visual learning, which refers to the enhancement of visual skills after training or exposure^3–7^ and is widely utilized to study adult neural plasticity in the visual system^8–12^, changes in sensory coding (e.g., sharpened tuning curve^13–15^, increased response gain^16–18^) have been present during encoding. At the same time, growing evidence is showing that consolidation also plays a critical role in neural plasticity.

Consolidation transforms fragile memory traces into long-term storage, resulting in maintained or improved behavioral performance, often considered as an “offline” gain. Research examining various forms of visual learning, including fine feature discrimination^19–22^, visual detection^23–26^, facial expression discrimination^27^, and complex natural and man-made object recognition^28^, has consistently demonstrated the importance of consolidation in the effectiveness of learning.

Consolidation processes are not limited to the realm of sleep; they also occur during wakefulness. A vast majority of research has been dedicated to investigating sleep consolidation, elucidating the neural oscillatory patterns, sleep spindle activity, slow-wave characteristics, blood-oxygen-level-dependent (BOLD) responses, neurotransmitter dynamics, and the role of different sleep stages in consolidation^19,22,26,28–33^. In contrast, the understanding of wakeful consolidation remains elusive but has gained increasing attention^10,20,34^. Wakeful consolidation extends over a temporal window following initial learning, whose length depends on customized training^20,21,23^, and is characterized by unstable memory traces and neural reactivation. To date, only a handful of studies have assessed the trace instability: introduction of an interfering task following the conclusion of training disrupted wakeful consolidation and led to a subsequent deterioration in behavioral performance during a retest^20,35^. Regarding neural reactivation during wakeful consolidation, a few studies have reported enhanced BOLD signals and increased BOLD decoding accuracy in visual areas after training on simple or complex visual stimuli^27,36^. However, it is important to emphasize that available evidence on neural reactivation during wakeful consolidation is still scarce. It is worth noting that, two studies have investigated neuro-metabolites and observed changes in the ratio between the primary excitatory neurotransmitter glutamate and the primary inhibitory neurotransmitter γ-aminobutyric acid (GABA) in the visual cortex after training^37,38^. These observations suggest the existence of an "E/I ratio" that undergoes changes during wakeful consolidation, underscoring the necessity for further investigations at this offline stage.

The phenomenon of neural reactivation during wakeful consolidation holds promise as a potential avenue for manipulating and influencing newly acquired memory traces. Until recently, a few studies have been conducted to explore the effectiveness of interventions in modulating wakeful consolidation of visual learning. Repetitive and theta-burst transcranial magnetic stimulations (TMS), which were documented to suppress neural excitability^39–41^, disrupted the wakeful consolidation when applied after the completion of training^21,25^. In the meantime, anodal transcranial direct current stimulation (a-tDCS) has been reported to increase neural excitability^42–47^ and facilitate wakeful consolidation^48–50^. These findings underscore the potential of modulating wakeful consolidation by manipulating neural excitability for targeted enhancement of specific functions within the visual system.

Repetitive visual stimulation (RVS) at different frequencies (delta^51,52^, theta^53^, alpha^54–57^, beta^51,58^, and gamma^59–63^) has emerged as a promising tool for modulating cognitive functions, neuro-electrophysiology, and cerebral hemodynamics, and even ameliorating Alzheimer’s disease^60,61^. Among them, alpha-frequency (8-12 Hz) RVS has garnered substantial evidence for its effects in enhancing early ERP (event-related potential) components^64–80^ and BOLD signals in the visual cortex^80,81^. In RVS studies, these effects are commonly acknowledged as non-invasive markers of long-term potentiation-like (LTP-like) plasticity^82^ and are thought to be associated with glutamatergic and GABAergic activities^64,69,81,83–85^, possibly also with neural excitability^64^.

Here, we attempt to investigate two unexplored but important questions: 1) can photic stimulation, in addition to behavioral methods and brain electrical/magnetic stimulations, influence wakeful consolidation, and 2) what are the neural underpinnings giving rise to the modulations? To answer these questions, we adopted the RVS paradigm and investigated its impact on wakeful consolidation in five experiments. Participants were exposed to different frequencies of RVS immediately after completion of visual training, and we collected neuroimaging data of multiple modalities. Our study revealed previously unknown and frequency-dependent impacts of RVS on wakeful consolidation of visual learning, as well as on ERPs and neurotransmitter concentrations.

## Results

### RVS modulates wakeful consolidation in a frequency-dependent manner

Our primary objective was to examine the potential of alpha-frequency photic stimulation, which had been proposed to act as a tetanizer on the human visual cortex and induce LTP-like plasticity^65,67,78^ (for review, see^79,86^), to enhance visual ability through strengthening wakeful consolidation. Meanwhile, 1-Hz RVS was found to weaken visual performance^87,88^.

Therefore, we investigated the frequency effect of RVS^89^ by adopting three different frequency conditions: 10, 1, and 0 Hz (served as the control condition). Here in Experiment 1, we utilized an orientation discrimination task (ODT) as the training and test paradigm of visual learning^90^, where two embedded-in-noise gabors with slightly different orientations appeared successively in either the lower left or right quadrant of the visual field and participants discriminated the rotation direction, either clockwise or counterclockwise (Fig. 1a, see ‘Orientation discrimination task’ in Methods).

**Fig 1.**
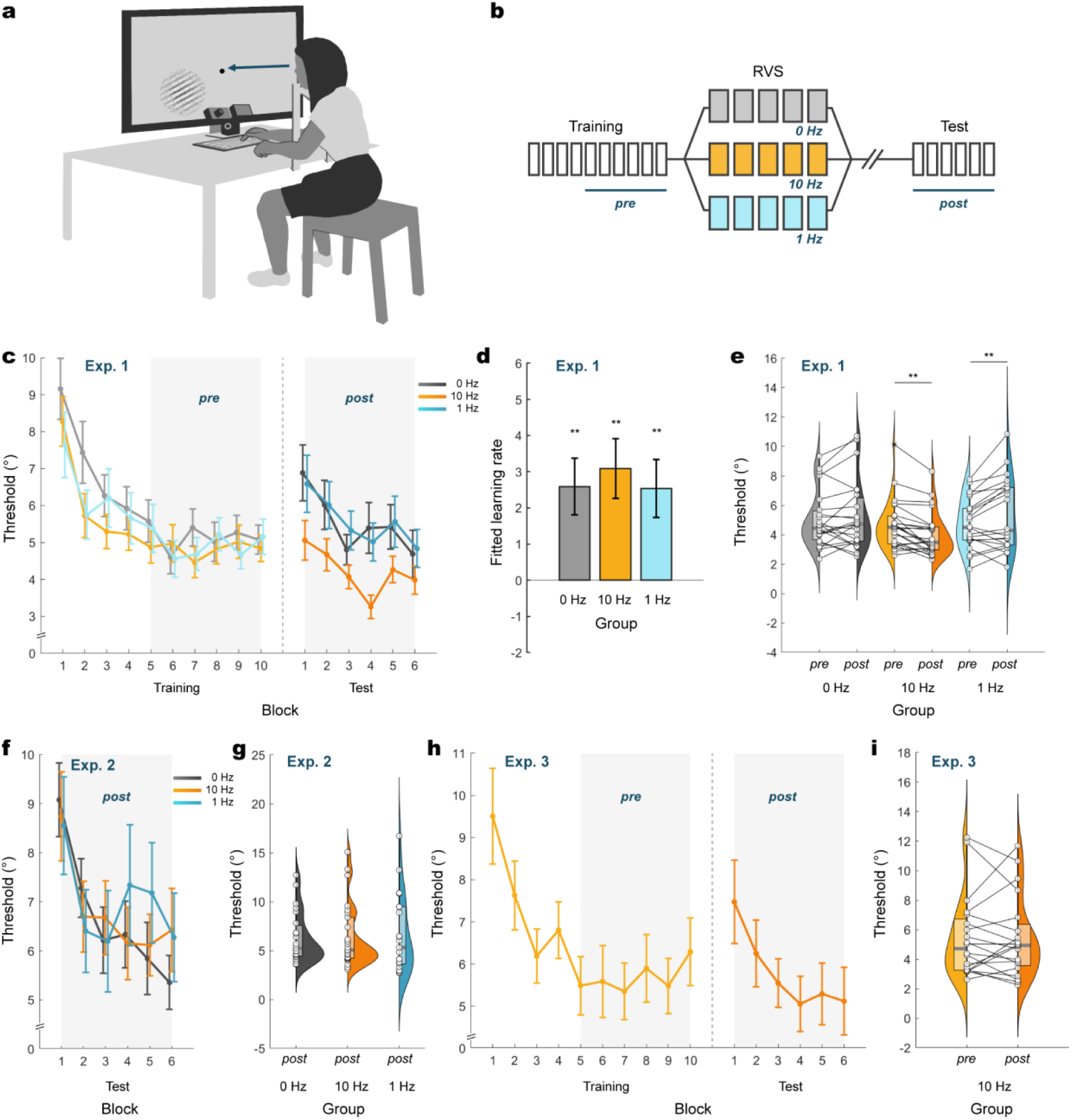
Experiments 1 to 3. **a**, Setup of behavioral experiments. Participants sat with their heads immobilized by a chin rest and fixated on the central dot on the screen. In the ODT, participants had to discriminate the orientation of a peripheral gabor and give their response using a keyboard. Eye movements were monitored using an eye tracker placed in front of the participants. **b**, Experimental procedure. Participants underwent the ODT training, RVS, rest, and the ODT test. The three RVS conditions were 0, 10, and 1 Hz. **c**, Learning curves in Experiment 1. Participants from all three groups exhibited training effects, indicated by the descending discrimination thresholds. A group difference in discrimination threshold was evident during the test. Shaded areas denoted data included in the ANOVA. **d**, All three groups demonstrated improvement during the training, indicated by above-zero learning rates. **e**, Discrimination thresholds in Experiment 1. Discrimination thresholds decreased after the 10-Hz RVS, increased after the 1-Hz RVS, and had no significant change after the 0-Hz RVS. Open circles, filled curved areas, transparent squares, and the horizontal lines within the squares represent individual participants, scaled probability density estimates, the 25-75% quantiles, and medians. **f**, Discrimination thresholds from the test of Experiment 2. **g**, No between-group difference in discrimination threshold was found for “naïve” participants in Experiment 2. **h**, Discrimination threshold curves in Experiment 3. **i**, Ten-Hz RVS did not change the discrimination threshold in Experiment 3. Error bars are MSEs. ** p<0.01.

A total of 70 participants sat in a behavioral test room and underwent ODT training for 10 blocks (approximately 30 minutes), immediately after which they were exposed to 5 blocks (25 minutes) of high-SNR (signal-to-noise-ratio) and high-contrast gratings flickering at different frequencies, i.e., RVS. Participants were randomly assigned to three groups (0 Hz, N = 23; 10 Hz, N = 25; and 1 Hz, N = 22) (see ‘Repetitive visual stimulation’ in Methods). Following RVS, participants underwent a 1.5-hour sleep-free break before completing a test consisting of 6 ODT blocks (approximately 20 minutes). We conducted a repeated measure ANOVA with a session (pre- and post-RVS) × group (0, 10, and 1 Hz) design to analyze the discrimination thresholds (see ‘Behavioral measurements’ in Methods). The “pre” session was defined as the last 6 blocks in the training, while the “post” session was defined as the 6 blocks in the test (Fig. 1b).

We found that participants’ performance gradually improved during the training (Fig. 1c). We conducted a paired t-test between the 1^st^ and 10^th^ training blocks and found a significant reduction in discrimination threshold (*t*_(68)_ = 8.996, *p* < 0.001, Cohen’s *d* = 1.083). We then fitted individual participants’ data to a power function to obtain individual learning rates and found that every group had an above-zero learning rate (one-sample t-test, Bonferroni adjusted. 0-Hz: *t*_(22)_ = 3.288, adjusted *p* = 0.006, Cohen’s *d* = 0.686; 10-Hz: *t*_(24)_ = 3.75, adjusted *p* = 0.003, Cohen’s *d* = 0.750; 1-Hz: *t*_(21)_ = 3.185, adjusted *p* = 0.006, Cohen’s *d* = 0.680. Fig. 1d. See ‘Supplementary Information’ for details).

We then analyzed the effects of group and session on discrimination thresholds. The repeated measure ANOVA revealed a significant interaction between group and session (*F*_(2,67)_ = 10.772, *p* < 0.001, partial *η*^2^ = 0.243), while neither group (*F*_(2,67)_ = 1.368, *p* = 0.262, partial *η*^2^ = 0.039) or session (*F*_(1,56)_ = 0.353, *p* = 0.554, partial *η*^2^ = 0.005) showed a significant main effect.

We further conducted an analysis of the simple main effect of session and found that participants demonstrated performance improvement after 10-Hz RVS (*F*_(1,67)_ = 11.519, *p* = 0.001, partial *η*^2^ = 0.147), performance deterioration after 1-Hz RVS (*F*_(1,67)_ = 8.434, *p* = 0.005, partial *η*^2^ = 0.112), and unaffected performance after 0-Hz RVS (*F*_(1,67)_ = 1.713, *p* = 0.195, partial *η*^2^ = 0.025) (Fig. 1e). These results showed that flickers at 10 and 1 Hz exerted opposite effects on wakeful consolidation, consequently resulting in improvement or deterioration in visual performance.

### RVS alone fails to modulate visual performance

In Experiment 1, we demonstrated that the 10-Hz RVS boosted wakeful consolidation after visual training, while the 1-Hz RVS disrupted it. In Experiment 2, we sought to answer the question: does RVS have to be implemented upon wakeful consolidation to modulate visual performance or can RVS itself influence following visual performance? To test the dependency of RVS-induced effects on prior training, we conducted Experiment 2.

In Experiment 2, the procedure was the same as that in Experiment 1 except that the ODT training was removed. Seventy newly recruited participants were randomly assigned to the three groups: 0-Hz (N = 23), 10-Hz (N = 26), and 1-Hz (N = 21). The participants were naïve to ODT when exposed to 25 minutes of RVS, after which they had a sleep-free break for 1.5 hours and then underwent a 20-minute test on ODT. A one-way ANOVA with group as the between-subject factor showed no group difference in discrimination threshold during the test (*F*_(2,67)_ = 0.061, *p* = 0.940, partial *η*^2^ = 0.002; Fig. 1f,g). For a more rigorous comparison, we fitted individual data in the test with a power function and compared the learning rates between groups. Results showed no between-group difference (*F*_(2,67)_ = 0.208, *p* = 0.813, partial *η*^2^ = 0.006) and that all three groups demonstrated moderate improvement (one-sample t-test on learning rate against zero, Bonferroni adjusted. 0-Hz: *t*_(22)_ = 2.652, adjusted *p* = 0.045, Cohen’s *d* = 0.553; 10-Hz: *t*_(25)_ = 1.939, adjusted *p* = 0.060, Cohen’s *d* = 0.380; 1-Hz: *t*_(20)_ = 2.338, adjusted *p* = 0.060, Cohen’s *d* = 0.510. see ‘Supplementary Information’ for details). These results demonstrated that the RVS of 10 or 1 Hz *per se* cannot exert influence on visual performance and needs to be implemented after training to take effect.

### Ten-Hz flickering uniform disk yields no effect on wakeful consolidation

A possible explanation for the facilitation effect of the 10-Hz flicker, as we demonstrated in Experiment 1, is that the 10-Hz flicker might have entrained alpha oscillation in the visual cortex, which could therefore, for example, enhance attention during ODT^90,91^. In Experiment 3, we tested whether the 10-Hz RVS effect can be accounted for by possible entrainment of neural oscillatory activity at alpha frequency or whether RVS has to contain orientation information to facilitate the wakeful consolidation of ODT.

Here we changed the RVS stimuli from oriented gratings to black and white uniform- luminance round disks, which have been documented to entrain alpha oscillation in the visual cortex^56,92^. The disk flickers had the same size, the same average luminance, and the same temporal contrast as the grating flickers in Experiments 1 and 2. The procedure here was identical to that of Experiment 1: participants underwent the training, RVS, a 1.5-hour break, and the test. We found that the 10-Hz disk flicker failed to reduce the discrimination threshold (N = 20, paired t-test: *t*(19) = 0.052, *p* = 0.959, Cohen’s *d* = 0.012; Fig. 1h,i). This suggests that the 10-Hz RVS effect cannot be explained by only alpha entrainment and that RVS has to contain orientation information to facilitate the wakeful consolidation.

### Ten-Hz RVS strengthens the ERPs associated with ODT

Previous studies of RVS without visual training have demonstrated that flickers with a frequency adjacent to 10 Hz induced LTP-like changes in ERPs^65–78^. In these studies, ERPs were measured during passive viewing (e.g. Teyler and colleagues^65^). However, it remains unknown whether the 10-Hz RVS, which we previously showed to improve visual task performance, also induces changes in task-associated ERPs. Therefore, in Experiment 4, we investigated whether the behavioral improvement induced by the 10-Hz grating RVS was accompanied by strengthened ERPs (i.e., LTP-like plasticity) evoked by the stimuli in ODT.

In Experiment 4, we recruited 45 participants and assigned them to the 0-Hz (N = 15), 10-Hz (N = 15), and 1-Hz (N = 15) groups. We recorded electroencephalogram (EEG) throughout the training, RVS, and the test. Here, the procedure was the same as that in Experiment 1 except for a longer break of 4 hours between RVS and the test. Careful optimization of stimulus parameters was also undertaken in this experiment to enhance the discernability and reliability of ERPs, given the dependence of ERPs on the physical characteristics of visual stimuli^93–95^ (see Table 1 for details and ‘Orientation discrimination task’ in Methods).

**Table 1.**
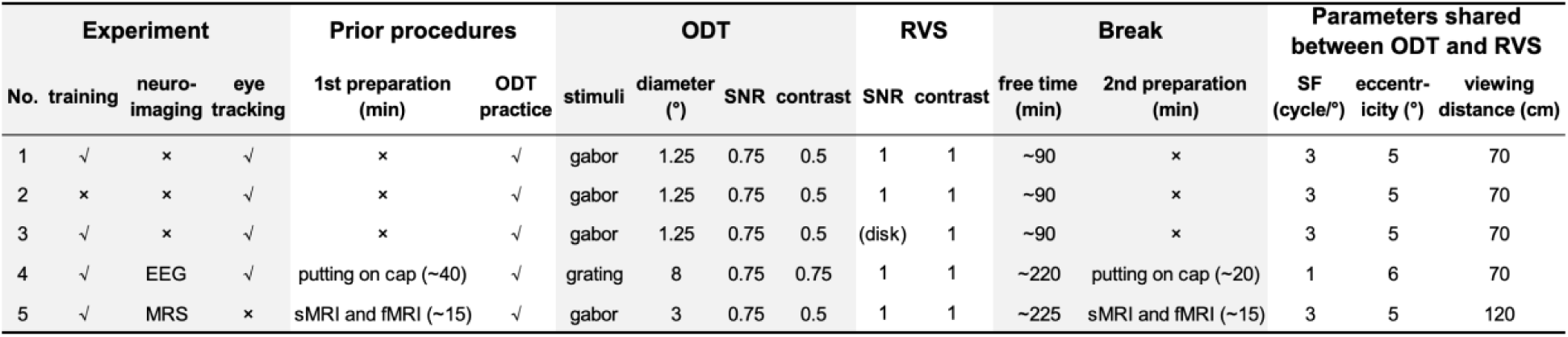
Differences in experimental procedure and stimulus design across experiments. SF is spatial frequency and SNR means the signal-to-noise ratio. Eccentricity is the distance from central fixation to the stimulus center. Acquisition of structural data is denoted by sMRI (structural magnetic resonance imaging). The parameters in Experiment 4 (the EEG experiment) were chosen to maximize the induction of ERPs.

To obtain the ERPs, we extracted EEG epochs around the onset of the first grating in each ODT trial with a duration of 350 ms, which were then baseline-corrected. These epochs were extracted from electrodes P5 and P6 of the 10-20 international EEG system, which were commonly selected for capturing ERPs elicited by peripheral stimuli^96–98^. P5 and P6 were labeled as “ipsilateral” or “contralateral” according to the parafoveal region (lower left or right quadrant) where the grating was presented (Fig. 2a,b; see ‘EEG’ in Methods).

**Fig 2.**
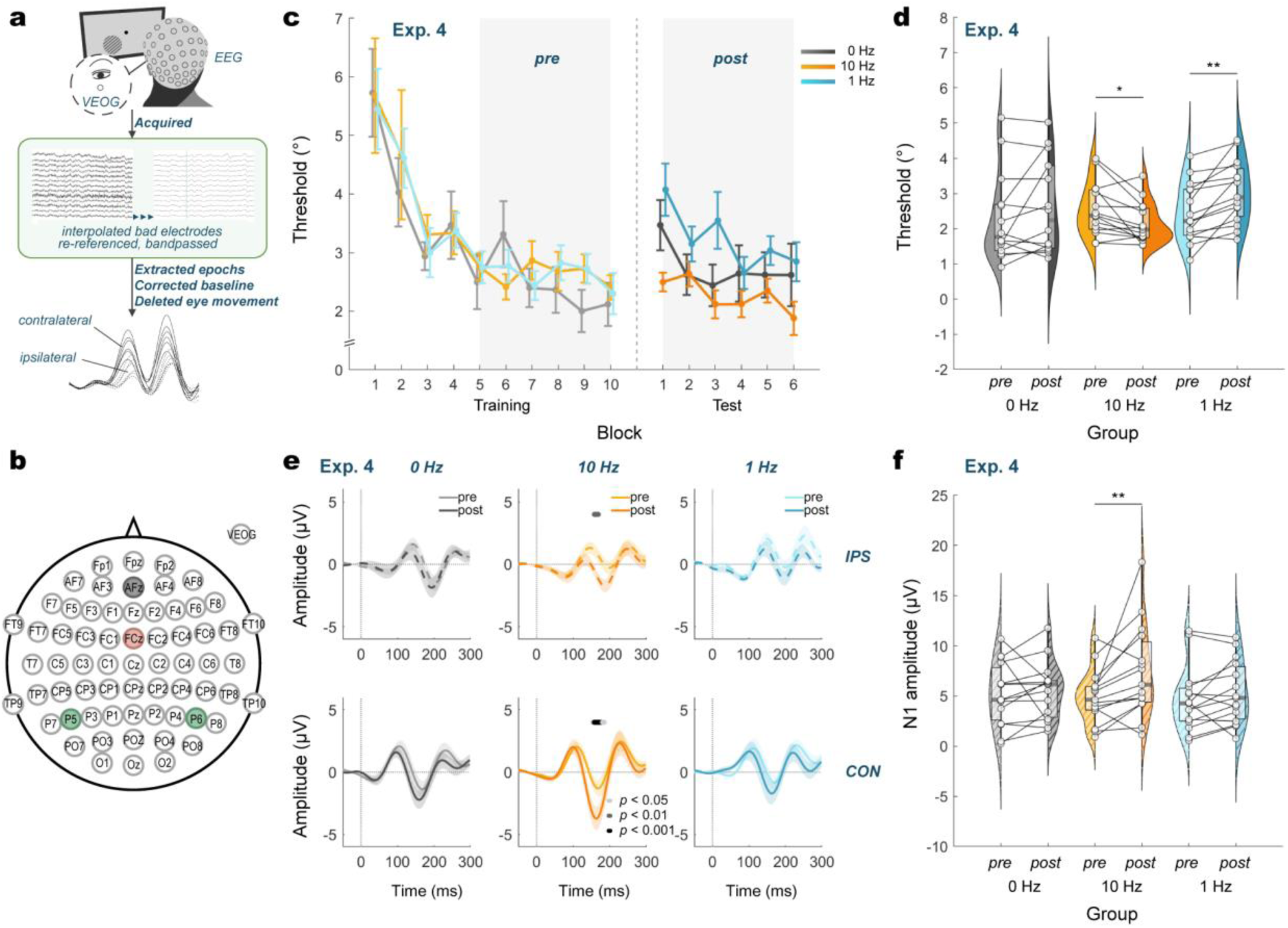
Experiment 4: 10-Hz RVS strengthens the ERPs. **a**, Setup of the EEG experiment. EEG and vertical electrooculogram (VEOG) signals were recorded during ODT and were preprocessed. Depicted were data from a representative participant. Dashed and solid curves are ERPs from electrodes on the ipsilateral and contralateral hemispheres, respectively. **b**, Electrode sites on EEG cap. A 64-channel EEG cap was used. EEG signals were referenced at FCz (pink) and grounded at AFz (gray). P5 and P6 were selected for ERP analysis (green). **c**, Learning curves in Experiment 4. **d**, Discrimination thresholds decreased after the 10-Hz RVS, increased after the 1-Hz RVS, and had no significant change after the 0-Hz RVS. Open circles are individual participants. Filled curved areas are scaled probability density estimates and transparent squares are the 25-75% quantiles. Thick gray lines within the squares are medians. **e**, ERPs under 12 combinations (2 hemispheres × 3 groups × 2 sessions). Upper and lower panels are ERPs from the ipsilateral (IPS) and contralateral (CON) electrodes. Shaded areas are MSEs. Vertical lines at 0 ms represent stimulus onset. ERPs after the 10-Hz RVS exhibited a strengthened negative peak at around 160 or 190 ms at the CON and IPS electrodes. **f**, N1 amplitudes. The amplitude of the N1 component was enhanced only by the 10-Hz RVS. Error bars are MSEs. * p<0.05, ** p<0.01.

Experiment 4 replicated the behavioral findings in Experiment 1. The effect of session (pre and post) was not significant (*F*_(1,41)_ = 3.000, *p* = 0.091, partial *η*^2^ = 0.068), nor was the effect of group (*F*_(2,41)_ = 0.774, *p* = 0.468, partial *η*^2^ = 0.036). A significant interaction between session and group was observed (*F*_(2,41)_ = 9.685, *p* < 0.001, partial *η*^2^ = 0.321). Further examination of the simple main effect of session with Bonferroni adjustment revealed that RVS modulated the wakeful consolidation of visual learning in different directions in a frequency-dependent manner: participants exhibited decreased, increased, and unchanged discrimination thresholds after the 10-Hz (*F*_(1,41)_ = 6.036, *p* = 0.018, partial *η*^2^ = 0.128), 1-Hz (*F*_(1,41)_ = 12.790, *p* = 0.001, partial *η*^2^ = 0.238), and 0-Hz (*F*_(1,41)_ = 3.426, *p* = 0.071, partial *η*^2^ = 0.077) RVS, respectively (Fig. 2c,d). These results further validate the effectiveness of the 10-Hz and 1-Hz RVS that we demonstrated in Experiment 1, despite slight modifications in stimulus feature and experimental procedure. These results provide evidence that the RVS effects on wakeful consolidation were relatively long-lasting and persisted till even 4 hours post-RVS.

Next, we analyzed the ERP data. The ERP waveforms obtained from the ipsilateral and contralateral sites before and after RVS were shown in Fig. 2e, and an enhanced N1 component of the ERPs following the 10-Hz RVS was observed. That is, the ERP showed a more negative response in the para-N1 time range (Bonferroni adjusted) at both the contralateral and the ipsilateral sites for the 10-Hz group. No such effect was observed for the 0- and 1-Hz groups. In addition, we calculated the peak-to-peak amplitude of the N1 component extracted from the contralateral sites as suggested by Markand^99^ and conducted a repeated measure ANOVA with a session (pre and post) × group (0, 10, and 1 Hz) design (Fig. 2f). We found a significant main effect of session (*F*_(1,42)_ = 7.747, *p* = 0.008, partial *η*^2^ = 0.156) and a marginally significant interaction between session and group (*F*_(2,42)_ = 2.682, *p* = 0.080, partial *η*^2^ = 0.113). No effect of group was observed (*F*_(2,42)_ = 0.609, *p* = 0.549, partial *η*^2^ = 0.028). Further analysis of simple main effects revealed that only the RVS at 10 Hz increased the amplitude of the N1 component (*F*_(1,42)_ = 12.063, *p* = 0.001, partial *η*^2^ = 0.223) but not that at 1 Hz(*F*_(1,42)_ = 0.879, *p* = 0.354, partial *η*^2^ = 0.021) or 0 Hz (*F*_(1,42)_ = 0.168, *p* = 0.684, partial *η*^2^ = 0.004). For the ipsilateral site, on the contrary, ANOVA yielded null effects (see ‘Supplementary Information’). These findings demonstrate that the 10-Hz RVS led to an augmented ERP component associated with ODT. The ERP effect might be an important factor contributing to the observed behavioral improvement and suggests an involvement of LTP-like plasticity.

### Ten-Hz RVS shifts the excitatory-inhibitory balance in the visual cortex towards excitation

The facilitation effect of the 10-Hz RVS was psychophysically assessed, and strengthened task-related neuro-electrical activity was indicated by the enhanced ERP component. Nonetheless, we still lack knowledge regarding the neurotransmitters underlying the development of the facilitation effect. Animal studies have well-documented the fundamental role of the excitatory neurotransmitter glutamate and the inhibitory neurotransmitter GABA in visual plasticity^64,100–103^. Recent studies have also demonstrated the importance of glutamatergic and GABAergic neural activities in the neural plasticity of the human visual system^69,81,84^. In fact, a delicate balance of excitation and inhibition, known as the glutamate/GABA equilibrium, is required for cortical plasticity to take place^104–106^. In order to gain a deeper understanding of the neural underpinnings of the RVS effects we demonstrated, we used proton magnetic resonance spectroscopy (MRS) to measure the changes in the relative concentration of glutamate and GABA during the training and the test.

In Experiment 5, 45 participants were evenly allocated to three groups (0, 10, and 1 Hz, each has N = 15). They were trained on ODT for about 30 minutes, exposed to 25 minutes of RVS, given a 4-hour sleep-free break, and then tested on ODT for about 20 minutes. While the overall setup was similar to that in Experiments 1, 3, and 4, visual stimuli were adapted to better fit the requirements of MRS scanning (see ‘Procedures and stimuli’ in Methods). MRS was acquired using MEGA PRESS^107,108^, and scanning spanned both the training and the test. The field of view (FOV) of MRS was 2 × 2.5 × 2.5 cm^3^, defined by an ROI (region of interest) localizer using functional magnetic resonance imaging (fMRI) to determine the area in the visual cortex responding to the peripheral gabor stimuli in ODT (see ‘fMRI and MRS’ in Methods).

The excitatory-inhibitory neurotransmitter balance, referred to as the E/I ratio, was examined as the ratio of glutamate and glutamine (Glx: Glu and Gln) to GABA within each participant’s FOV (Fig. 3a,b).

**Fig 3.**
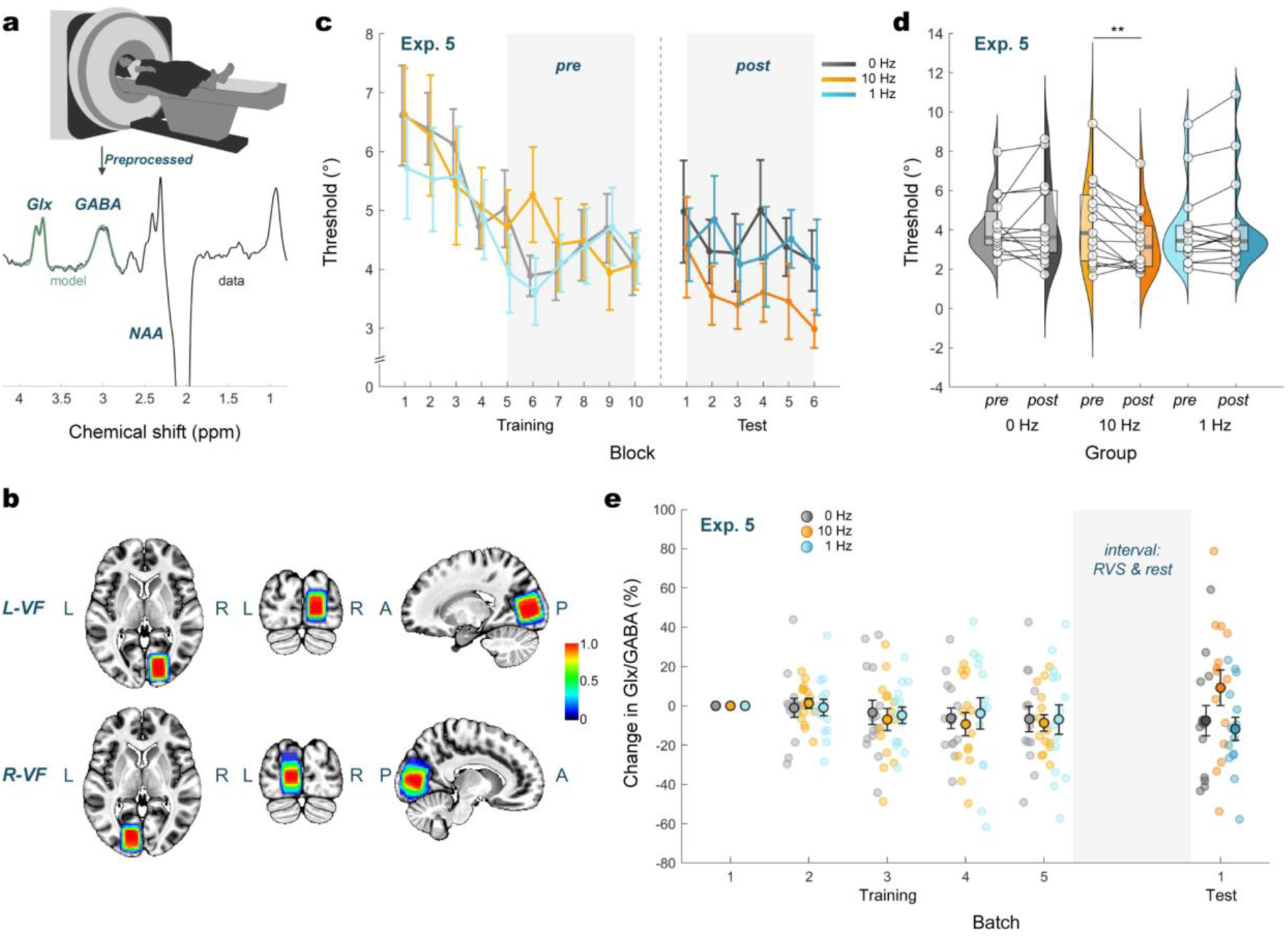
Experiment 5: ten-Hz RVS increases neural excitation in the visual cortex. **a**, Setup of the MRS experiment. Participants viewed the screen through mirror reflection from inside the scanner. MRS data were acquired during ODT and were preprocessed to obtain individual spectra. X axis is the degree of chemical shift expressed in part-per-million (ppm). **b**, FOVs employed during MRS acquisition. FOVs were selected based on the ROI localizer to cover the cortical areas activated by ODT in the early visual cortex. Participants were randomly assigned to be trained in the lower left or right visual field (VF). Legend color is the probability of a voxel being included in FOVs in the experiment. **c**, Learning curves in Experiment 5. **d**, Ten-Hz RVS decreased the discrimination threshold while null effects were found for both 1- and 0-Hz RVS groups. Open circles and filled curved areas are individuals and the scaled probability densities. Transparent squares are the 25-75% quantiles and the horizontal lines within indicate the medians. **e**, Batch analysis of E/I ratios. Ratios derived from each batch were normalized to the first batch. The E/I balance shifted towards inhibition during the training process. The shaded area represents the interval between the training and test, i.e., RVS and the rest. Gray, yellow, and blue dots with black edges denote the group average changes in the E/I ratio of the 0-, 10-, and 1-Hz RVS groups, respectively. Error bars are MSEs. ** p<0.01.

Similar to Experiments 1 and 4, participants in Experiment 5 also showed a performance gain in ODT after the 10-Hz RVS, despite slight stimulus differences. A significant interaction between group and session was found (*F*_(2,42)_ = 5.852, *p* = 0.006, partial *η*^2^ = 0.218), but there was no main effect for either group (*F*_(2,42)_ = 0.173, *p* = 0.842, partial *η*^2^ = 0.008) or session (*F*_(1,42)_ = 1.217, *p* = 0.276, partial *η*^2^ = 0.028). Examination of simple main effects of session showed significance for the 10-Hz group (*F*_(1,42)_ = 11.753, *p* = 0.001, partial *η*^2^ = 0.219), but not the 1-Hz (*F*_(1,42)_ = 0.441, *p* = 0.510, partial *η*^2^ = 0.010) or the 0-Hz (*F*_(1,42)_ = 0.728, *p* = 0.399, partial *η*^2^ = 0.017) group (Fig. 3c,d). One possible explanation for the lack of deterioration in ODT performance after exposure to the 1-Hz RVS is the changes in the experimental environment (noise, vibration, nervousness, etc.). Participants’ performance during pre (i.e., last 6 training blocks) was at a peri-floor level, making further impairment less likely than in Experiments 1 and 4.

With the acquired MR data, we calculated the average distance of individual FOVs from the mean Montreal Neurological Institute (MNI) coordinates. For participants who were trained in the lower left quadrant, FOV was in the right hemisphere (mean ± MSE: x = 16.96 ± 0.24 mm, y = −81.17 ± 0.56 mm, z = 8.00 ± 0.61 mm). For those who received training in the lower right quadrant, FOV was in the left hemisphere (mean ± MSE: x = −13.39 ± 0.28 mm, y = −83.77 ± 0.43 mm, z = 6.55 ± 1.02 mm). Small MSEs indicate that the individual difference in FOV location was small.

Analysis of MRS data obtained within the FOV revealed that training on ODT, which demands fine discrimination, decreased the E/I ratio in the visual cortex. This is consistent with Frangou and colleagues’ finding^109^. Similar to their study, we constructed a linear mixed effects (LME) model. We also analyzed the E/I ratio using a sliding window (batch) with a window length of 6 blocks (approximately 12 minutes) and with a step length of 1 block during the training, normalized to the first batch. The LME model showed that, before RVS, participants from each group had comparable E/I levels in the visual cortex (*F*_(2,114)_ = 0.114, *p* = 0.893).

Interestingly, the E/I ratio decreased during the training (*F*_(1,114)_ = 9.385, *p* = 0.003; Fig. 3e). This suggests that training on fine orientation discrimination induced a decrease in visual cortical excitation, accompanying the training-induced behavioral improvement.

Most importantly, we found that the 10-Hz RVS, previously found to strengthen the task-related ERPs, shifted the E/I balance towards the excitatory side when implemented immediately after the training. We used a second LME model (group × session) to analyze the normalized E/I ratio of each group in each session. The sessions here are pre (the last batch, i.e., last 6 blocks, in the training) and post (the one batch, i.e., 6 blocks, in the test). Bonferroni adjusted simple main effects showed a significant increase in the E/I ratio after the 10-Hz RVS (*F*_(1,37.966)_ = 4.851, *p* = 0.034) but not the 1-Hz (*F*_(1,37.966)_ = 0.365, *p* = 0.549) or 0-Hz RVS (*F*_(1,39.358)_ = 0.017, *p* = 0.897). The results demonstrate that the 10-Hz RVS elevated neural excitation in the visual cortex, accompanying the 10-Hz RVS-induced behavioral improvement.

The results here suggest that the plasticity resulting from the training and the plasticity induced by the 10-Hz RVS might operate through different neurochemical mechanisms.

Specifically, continuous training on feature discrimination is associated with a decreased visual cortical excitation, as opposed to the plasticity induced by the 10-Hz RVS, which is associated with an increased visual cortical excitation.

## Discussion

In this study, we reported a new method, i.e., the 10-Hz photic stimulation, to facilitate the offline processing of encoded sensory information. Our investigation not only emphasized the significance of wakeful consolidation beyond online processes and sleep, but also delved into the neural substrates underlying the effect of the 10-Hz RVS. In Experiment 1, we showed that, after visual training, the 10-Hz RVS facilitated wakeful consolidation, while the 1-Hz RVS impaired it. In Experiment 2, we showed that the RVS effects could not be elicited by RVS *per se*; instead, RVS needed to be implemented at the post-training stage, i.e., wakeful consolidation. In Experiment 3, we showed that the 10-Hz facilitation effect was not simply a result of entrainment of alpha oscillation. In Experiment 4, we analyzed task-associated ERPs and found an augmented N1 component after the 10-Hz RVS, indicating LTP-like plasticity in task-related visual functions. In Experiment 5, we investigated the glutamatergic and GABAergic activities underpinning the 10-Hz RVS effect and found an increased E/I ratio. In addition, our study is, to the best of our knowledge, the first to provide evidence on RVS-induced changes in neurotransmitter concentrations in humans.

Our study unprecedentedly demonstrated the potential of photic stimulation to be harnessed to modulate wakeful consolidation of visual learning. Photic stimulation offers significant advantages over traditional methods such as electrical or magnetic stimulation and drug intervention^21,24,25,48–50,110^. Photic stimulation provides practicality, user-friendliness, and broad applicability, thereby increasing its feasibility and acceptance in various research and practical contexts. Furthermore, photic stimulation can be meticulously tailored to selectively target specific neurons, circuits, and pathways in the visual system^111^, which can be helpful in elucidating intricate neural mechanisms underlying diverse aspects of visual functions.

Our study also demonstrated the essentiality of the wakeful consolidation window, despite the commonly held belief that the inattentive offline stage during daytime wakefulness is unproductive^112^. Moving beyond visual learning, a growing body of research in areas such as auditory learning, procedural memory, and declarative memory is also corroborating the significance of this critical offline window, wherein interventions can exert mnemonic influences^113–120^. However, once the window closes, the retention of learned information stabilizes, rendering it much less amenable to interventions^20^.

In this study, we demonstrated facilitated wakeful consolidation by the 10-Hz RVS and that this was accompanied by an augmented N1 component. This finding is consistent with previous research^65–68,70–72,74,76–78^ despite variations in stimulus design. The N1 complex was proposed to be associated with visual discrimination rather than detection^121,122^. The observed increased negativity of N1 could potentially be attributed to LTP-like plasticity within discrimination-associated neural substrates.

In addition to changes in field potential activities (i.e., ERPs), our study revealed increased neural excitation in the visual cortex after the 10-Hz RVS. This link between neural excitation and the behavioral benefit aligns with findings from transcranial magnetic/electric stimulation studies on discrimination tasks. Decreasing neural excitation disrupts, while increasing it facilitates, the wakeful consolidation of visual learning. For example, both 1-Hz repetitive TMS (rTMS) and continuous theta burst stimulation (cTBS) reduce cortical excitability^39–41,123^, impairing wakeful consoldiation^21,25^. Conversely, a-tDCS enhances neural excitation in the visual cortex^42–47,124^ and improves wakeful consolidation^48,49^. However, it is worth noting that studies adopting a detection task yielded results opposite to those with a feature discrimination task. Peters and colleagues reported impaired wakeful consolidation by a-tDCS after training on a detection task^24^. Since fine visual feature discrimination- and detection-based training shift the E/I balance towards inhibition and excitation, respectively^109^, adjusting the neural excitation level towards equilibrium during wakeful consolidation seemed to benefit the offline processing of stored information. However, further investigation is required to determine the effects of inhibitory intervention on detection tasks before formulating a conclusive proposition.

In the current study, we showed that the uniform disk flicker, though able to entrain alpha oscillation in the visual cortex^56,92^, failed to facilitate wakeful consolidation. An intriguing question would be: why is the effect specific to alpha frequencies if it is not a result of alpha entrainment? We observed a transient increase in alpha power in the occipital lobe by the 10-Hz grating RVS in Experiment 4 (the EEG experiment, Fig. S2) but no alpha entrainment (Fig. S4). This is consistent with alpha-frequency transcranial alternating current stimulation (tACS) research indicating that brief epochs of stimulation failed to entrain alpha oscillations but only resulted in a transient increase in alpha power^55^. Furthermore, they propose spike-timing-dependent plasticity (STDP) involvement during the transient power increase, potentially underpinning the observed LTP-like plasticity. The plasticity is specific to 10 Hz probably because of the intrinsicity and dominance of alpha frequency in the visual cortex^55,125–127^, akin to the beta band’s significance in auditory processing^128–131^ and its association with auditory LTP-like plasticity^132–134^.

We also reported that the RVS-induced plasticity was dependent on prior training, as evidenced by the lack of impact on naïve participants. This complements Marzoll’s discovery that proficient, well-trained participants experienced performance deterioration following 10-Hz RVS^135^. Taken together, well-trained, newly trained, and naïve participants represent different learning stages: well-trained participants with saturated performance and stabilized neuronal substrates^136^ may possess different neurochemical attributes in their visual system compared to the newly trained. In contrast, cortices naïve to training lack the biological milieu necessary for facilitation to take place.

Interestingly, we unraveled two forms of macroscopic neural plasticity in the visual system: training-induced plasticity and tetanization-induced plasticity. These two forms of plasticity relied on diametrically opposite changes in neurotransmitter concentration despite having consistent beneficial effects on behavioral performance. We observed a reduction in neural excitation during the training. Here are two possible explanations. First, task repetition may lead to visual adaptation and overlearning, both linked to reduced cortical excitation^37,137^. Second, discrimination tasks decrease visual cortical excitation. Here we propose a hypothetical framework to integrate cortical excitatory dynamics during training and tetanization. Under this framework, task type influences neurotransmitter dynamics during training: discrimination training sharpens feature representation supported by parvalbumin-positive GABA interneurons^138^ and increases GABA concentration^109^, while conversely, detection training reduces GABA concentration^109^, increases glutamate concentration^38^, and increases the E/I ratio^37,38^ in visual cortical areas (Fig. 4). Meanwhile, it is also noteworthy that the SNR of 3.0 T MRS imposes constraints on the length of the sliding window. In our study, the fixed window length (6 blocks) might be too coarse to capture non-monotonic changes in the E/I ratio, should they exist.

**Fig 4.**
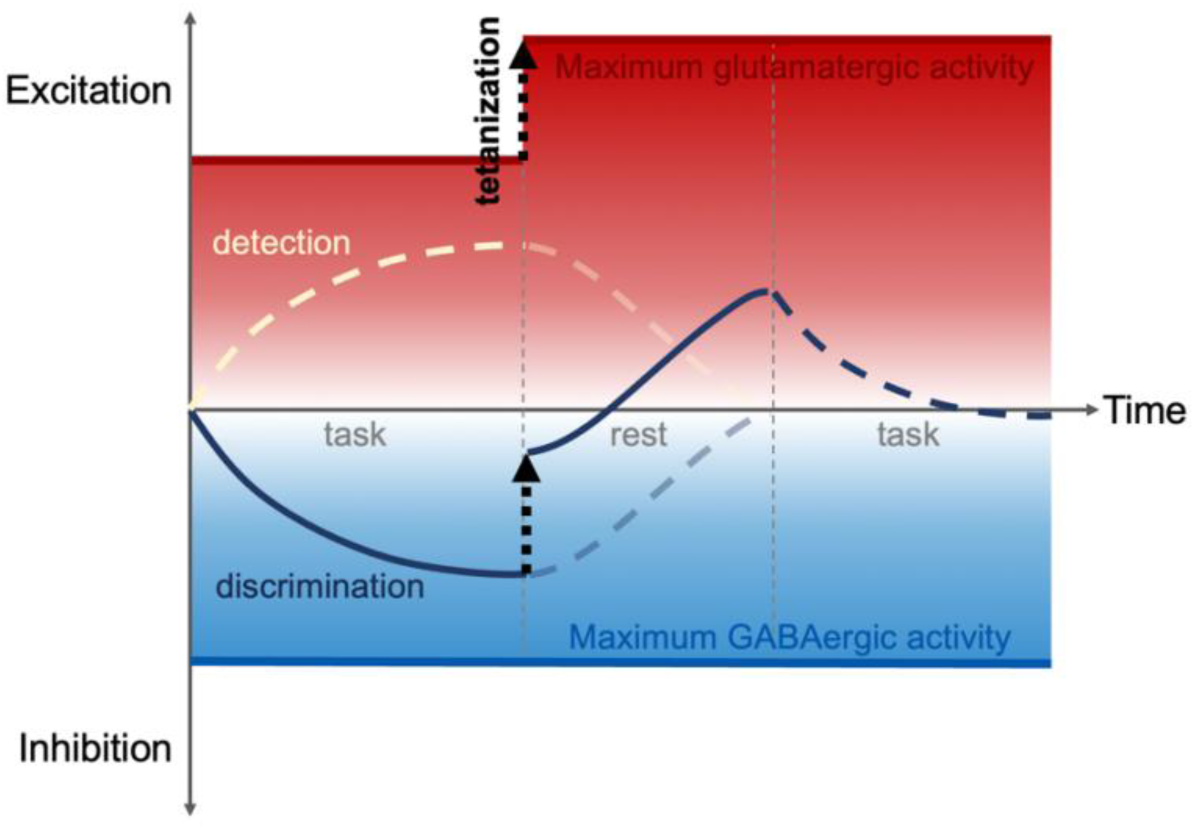
Framework of neurotransmitter dynamics during training and tetanization. E/I ratio is reduced during training on a fine-feature discrimination task but is elevated after visual tetanization. Our explanation is that 10-Hz RVS induces LTP-like plasticity in glutamatergic activities, shifting the overall process towards excitation and resulting in both augmented field potentials and improved behavioral performance. Beige and blue curves indicate discrimination and detection tasks, respectively. Solid curves represent observed data in the current study while dashed curves represent hypothesized changes or conclusions from other studies.

On the other hand, the 10-Hz RVS induced an increase in neural excitation. This finding is in line with previous studies showing that LTP-like plasticity is not only dependent on glutamatergic transmission in the visual cortex, but also increases glutamate signaling. Blockage of glutamatergic activity by competitive antagonist of glutamate receptors abolishes the augmentation of ERPs^64^, while enhancement of glutamatergic signaling by receptor agonist amplifies the LTP-like effect^69^. In human studies, the effect of alpha RVS depends on the level of pre-stimulation glutamate concentration^81^. Alpha RVS, in return, increased glutamate receptor expression^85,139^. Hence, it is reasonable to hypothesize that an increase in glutamatergic neuronal activity serves as one of the primary mechanisms driving the observed LTP-like plasticity, constituting the tetanization part of the framework (Fig. 4). Nevertheless, it is important to note that MRS measures bulk concentration of neurotransmitters and cannot directly assess their specific origins^140^. Thus, future animal studies are warranted to delve further into this hypothesis.

In summary, we discovered a new way to modulate the wakeful consolidation of visual learning, while also investigating the neural substrates underlying this modulation. Furthermore, we proposed a framework which illustrates that task-related neural underpinnings, actively engaged after training^27,36–38^, provide an optimal stage for alpha RVS to induce glutamate-associated LTP-like plasticity. The present study contributes novel insights into the neural mechanisms underlying wakeful consolidation and offers new avenues for neuromodulation.

Future studies could explore alternative scenarios involving various visual tasks, RVS characteristics, timing strategies, etc., and the impact of tetanization on offline neural processes in other domains such as motor learning and semantic memory, thereby examining the validity of the falsifiable framework we provided.

## Methods

### Participants

A total of 250 participants met all screening criteria (see ‘Supplementary Information’) and provided written informed consent forms prior to experiment commencement. Among these participants, 70 (mean ± SD: 22.30 ± 3.12 years old, 44 female), 70 (21.38 ± 2.47 years old, 39 female), 20 (20.65 ± 2.46, 11 female), 45 (20.80 ± 2.15 years old, 14 female), and 45 (20.67 ± 1.94 years old, 24 female) participated in Experiments 1-5, respectively. All participants had normal or corrected-to-normal vision and no self-reported history or family history of neurological or mental disorders. Monetary compensation was provided to all participants for their involvement in the study. All experiments in this study were approved by the ethics committee of the School of Psychological and Cognitive Sciences, Peking University.

### Procedures and stimuli

Experiments 2-5 were derivatives from Experiment 1. In Experiment 1, the objective was to examine whether 10-Hz tetanization-mimicking RVS could enhance the early post-training consolidation of visual learning. Each participant first provided informed consent and filled out a screening questionnaire battery (see ‘Supplementary Information’). Only participants who met all screening criteria underwent the subsequent procedures, which consisted of ODT training of 10 blocks, RVS of 5 blocks, a sleep-free break of 1.5 hours, and finally, an ODT test of 6 blocks. Each training and test block comprised 50 trials, with a mandatory rest period of at least 50 seconds between blocks. In every RVS block, we presented 5-minute flickering stimuli at 10, 1, or 0 Hz, and participants rested for at least 20 seconds between RVS blocks. This experiment was conducted under a semi-double-blind condition where both the participants and a second experimenter who conducted the post-RVS test were unaware of the experimental condition (i.e., RVS frequency).

During the experiment, participants were seated in a dimly lit room and instructed to maintain their fixation on a central dot at all times during ODT and RVS. The viewing distance was about 70 cm. Eye movements were monitored using an eye tracker (Eyelink 1000 Plus, SR Research, Ontario, Canada). During the break between RVS and the test, participants were instructed to refrain from sleeping or engaging in physical or mental exercise and were required to wear a smart band (Honor Band 6, Honor Device, Shenzhen, China) on their wrist to monitor physical activity during the break.

Experiments 2-5 were conducted single-blindedly. In Experiment 2, the training was omitted. In Experiment 3, the gratings in RVS were changed into round uniform disks and only the 10-Hz group was included. In Experiment 4, the rest between RVS and the test was prolonged to 4 hours, and EEG was acquired throughout the training, RVS, and the test. In Experiment 5, we acquired structural and functional MRI data for the first time before the ODT training and a second time before the ODT test. MRS data were acquired during the training and test. The viewing distance was increased to about 120 cm in Experiment 5.

Visual stimuli used in ODT, RVS, and fMRI ROI localizer in all five experiments were generated using Matlab and Psychtoolbox-3^141^.

### Orientation discrimination task

The visual stimuli were presented on either a Display++ monitor (Cambridge Research Systems Ltd., Kent, UK, used in Experiments 1-4) or an MR-compatible BOLDscreen monitor (Cambridge Research Systems Ltd., Kent, UK, used in Experiment 5). Both screens had a refresh rate of 120 Hz and a resolution of 1920 × 1080 pixels, and were calibrated for gamma-correction to a linear 0-to-100 cd·m^-2^ before experiment commencement. A fixation dot with a diameter of 0.17° was constantly visible at the center of the screen against a gray background of half the maximum luminance of the monitor, i.e., 50 cd·m^-2^. ODT stimuli were either Gaussian-enveloped sinusoidal gabors (used in Experiments 1, 2, 3, and 5) or sinusoidal gratings without the envelope (used in Experiment 4), with orientations around either 26° or 154°, depending on the visual field where stimuli were presented. The stimulus parameters are detailed in Table 1.

In each trial, the first stimulus was displayed for 100 ms, followed by a 600-ms blank interval, and then a second 100-ms stimulus with a slightly altered orientation presented at the exact location of the first stimulus. Participants needed to indicate the direction of the rotation (clockwise or counterclockwise) by pressing one of two keys and were explicitly instructed not to rush their response. After the response was given, there was a lapse between trials, which lasted 500 ms in Experiments 1, 2, 3, and 5 but varied between 500 and 700 ms in Experiment 4 (i.e., the EEG experiment). Meanwhile, note that the length of between-stimulus intervals also jittered between 500 and 700 ms across trials in Experiment 4. This design was implemented to avoid EEG artifacts arising from the relatively close arrangement of stimuli and trials.

Each ODT block consisted of 50 trials, and the difficulty level (i.e., orientation difference) was controlled using a QUEST staircase procedure to estimate participants’ discrimination threshold at 75% accuracy. No feedback was provided during the ODT training or test. However, during the practice round, which took place before the training started, participants received feedback—the central fixation dot turned green after each correct response—in order to facilitate practice. The practice round consisted of 25 trials and employed orientations (116° or 64°) different from those in the ODT training and test (26° or 154°).

### Repetitive visual stimulation

RVS was administered immediately following the completion of the ODT training. The sinusoidal gratings used in RVS were noise-free and slightly larger in size than the gabors/gratings in the corresponding ODT, ensuring full coverage of the trained area. High similarity between the stimuli adopted in ODT and in RVS was ensured as sensory LTP-like plasticity is input-specific^66,67,79^. That is, gratings in RVS had the same spatial frequency and orientation as those in ODT. Detailed parameters can be found in Table 1.

Each RVS block lasted 5 minutes and we investigated the effects of RVS with different frequencies. In the 10-Hz RVS group, the grating intermittently flickered on and off with phase reversing at 10 Hz for 1 second after every 5 seconds of blank screen. Within each second of flickering, a grating of phase φ was visible for 16.67 ms, followed by an 83.33-ms blank period. This was then followed by a second grating of phase φ+180° for 16.67 ms and another 83.33-ms blank period. This sequence was repeated five times hence 1 second.

In the 1-Hz RVS group, the grating continuously flickered with phase reversing at 1 Hz for 5 minutes in each block. Specifically, after each time a grating of phase φ was visible for 16.67 ms, the screen went blank for 983.33 ms, followed by the appearance of a second grating of phase φ+180° for 16.67 ms and another 983.33-ms blank period. The design of the 10- and 1- Hz RVS groups is similar to that described in the study of Marzoll and colleagues^135^.

In the 0-Hz RVS group, 15 static gratings were presented, each with a duration of 1.67 seconds. They spread out over the entire 25 minutes of RVS. The intervals between every two gratings were randomized to eliminate frequency information while maintaining a comparable amount of visual stimulation with the other two groups.

We employed an oddball paradigm to ensure participants’ attention to the visual stimuli during RVS, which required participants to detect occasional changes in grating size. The oddball stimuli had a diameter of 0.5-0.75 times the original grating diameter. The spatial frequency of the oddball stimuli scaled with their grating size. After each block, participants were asked to indicate the number of smaller gratings they detected in the block using a numeric keyboard. Participants who answered incorrectly two times out of five were excluded from further analysis.

In Experiment 3 (i.e., the “uniform disk” experiment), gratings of phase φ and φ+180° were replaced with white and black uniform disks of the same diameter.

### fMRI localizer

The ROI localizer stimulus was presented on the BOLDscreen monitor, as mentioned earlier, and viewed from inside the MR scanner. The purpose of the localizer was to identify the areas in early visual cortex responding to the gabors in ODT. To achieve this, a black and white checkerboard pattern (gird size: 0.44°) was reversed at a frequency of around 4 Hz (3-5 Hz)^142^.

The choice of a variable reversal frequency allowed us to eliminate any potential interference from the 4-Hz visual stimulation on the effects of our specific frequencies of interest: 10, 1, and 0 Hz. The checkerboard subtended a circular area of 3°, with its center located 5° from the central fixation point. This location coincided precisely with the gabors in ODT in Experiment 5. The reversing checkerboard was presented for 15 seconds followed by a 15-second blank screen, constituting a cycle that was repeated 8 times, taking up a total of 4 minutes. During the localizer, participants were instructed to maintain their fixation on the central dot at all times.

### Data acquisition and analysis

#### Behavioral measurements

As for ODT, the geometric mean of the threshold values obtained from the last 6 blocks in the training was termed “pre” and was compared to the geometric mean of the threshold values from the 6 blocks in the test, which was termed “post.” Participants whose pre or post performance deviated by three or more standard deviations from the mean were excluded from further statistical analyses. We employed a 2×3 repeated measure ANOVA with a within-subject factor of session (pre and post) and a between-subject factor of group (0, 10, and 1 Hz) to analyze the ODT thresholds in Experiments 1, 4, and 5. In Experiment 2 (the “naïve” experiment), a one-way ANOVA (between-subject factor: group) was used for statistical analysis. In Experiment 3 (the “uniform disk” experiment), a paired t-test was used to compare pre and post performance for the 10-Hz RVS group.

#### EEG

Experiment 4 (the EEG experiment) was conducted in a magnetically shielded room, and data were collected using a 64-channel Ag/AgCl EEG cap (Easycap, BrainProducts, Gilching, Germany). EEG signals were continuously recorded at a sampling rate of 500 Hz with an FCz reference and preprocessed using EEGLAB (version 2021.1). EEG signals were re-referenced to bilateral mastoids and bandpass filtered (1-12 Hz^96^). Electrodes P5 and P6 were selected for epoch extraction because these electrodes exhibited the highest peak amplitude of the N1 component over their respective contralateral-to-stimulus hemisphere. Data from these two electrodes were then segmented into baseline-corrected epochs with a 50-ms pre-stimulus baseline and a 300-ms period following stimulus onset. To avoid neural response overlap between stimuli, only epochs corresponding to the first grating in a trial were included, as the ERPs induced by the second grating were visible before the ERPs induced by the first grating had returned to baseline^143^. Epochs with significant eye movement were removed (see ‘Supplementary Information’). ERPs from every participant were then averaged across the last two training blocks (pre) and the first two test blocks (post), according to the electrode (either P5 or P6) contralateral (CON) or ipsilateral (IPS) to the visual field in which the gratings were presented. Here, we included 2 blocks in each session for EEG data, as opposed to 6 blocks for behavioral data, due to the potential of visual training to enhance related ERP components^144^.

The contralateral hemisphere exhibited the N1 component before the ipsilateral hemisphere (Fig. 2a,e). Also, decreasing the stimulus size, increasing eccentricity from central fixation, and utilizing an LCD monitor instead of a CRT monitor might all contribute to the prolongation of ERP component latency^94,99,145,146^. N1 was hence defined as the negative peak occurring between 138 and 192 ms for the CON electrode and between 164 and 224 ms for the IPS electrode. The peak-to-peak amplitude of N1 at the CON electrode was calculated and analyzed using a 2×3 repeated measure ANOVA (within-subject: session; between-subject: group).

### fMRI and MRS

A 3-tesla Prisma MR scanner, a Syngo MR XA30 system (Siemens, Forchheim, Germany), and a head-neck 64 coil were used for acquiring structural and functional images and MRS. High-resolution T1-weighted structural images were obtained using the MPRAGE sequence with GRAPPA2 (TR 2300 ms, TE 2.98 ms, FOV 232 × 256 mm^2^, voxel size 1.0 × 1.0 × 1.0 mm^3^, 192 slices, and 5:03 minutes acquisition time). BOLD signals were collected using an echo-planar imaging sequence (TR 1500 ms, TE 30 ms, FOV 192 × 192 mm^2^, matrix 96 × 96, thickness 2 mm, 60 slices, and 4:09 minutes acquisition time).

BOLD signals were acquired during the ROI localizer using a block design. Two conditions (on and off, see ‘fMRI localizer’ in Methods for details), each with a duration of 15 seconds, were interleaved to allow the BOLD signal to return to baseline each time after the response to the checkerboard was elicited. Functional data were aligned to the first volume to correct for possible head motion during scanning and registered on individual structural data. We used a two-category (on and off) general linear model (GLM) and a paired t-test in SPM12^147^ to identify the activated areas during the ROI localizer. ROIs were defined as the activated areas alongside the calcarine fissure in the hemisphere contralateral to the localizer stimulus and were subsequently used in the determination of the FOV in MRS.

MRS was collected using the MEGA PRESS sequence^107,108^ provided by the University of Minnesota under a C2P agreement (TR 2000 ms, TE 68 ms, FOV 20 × 25 × 25 mm^3^, with VAPOR water suppression, variable acquisition time depending on task performance speed). Data processing was performed using a customized Gannet 3.1 toolbox^148^ in Matlab 2023a. We selected only the scans within every ODT block and discarded the scans entirely or partially outside the blocks. Scans were then reorganized into blocks so that each ODT block corresponded to one set of MRS scans. We then adopted a sliding window (referred to as “batch”) of the length of 6 blocks and the step size of 1 block, and calculated the average spectrum of each batch, yielding five spectra in the training and one spectrum in the test for each participant.

The concentrations of Glx and GABA were obtained by fitting the spectral lines corresponding to their respective positions. Data points with fitting errors exceeding 10% were excluded from further analyses, leaving 14 participants in each group. The E/I ratio was calculated as Glx to GABA, and the percentage change in E/I ratio relative to the first batch was further analyzed using LME models. The first LME model aimed to test whether the E/I ratio declined with training; therefore, it was designed as a change in E/I = repetitive covariate batch (1 to 5) × group (0, 10, and 1 Hz) with intercepts included. A second LME model with change in E/I = repetitive session (pre and post) × group (0, 10, and 1 Hz) with intercepts was used to examine the effects of both session and group.

For the visualization of FOVs, FOV geometries were transformed into a standard MNI coordinate system using individual structural data and registered onto an MNI152 standard brain image, as depicted in Fig. 3b.

## Acknowledgement

This study was supported by National Science and Technology Innovation 2030 Major Program (2022ZD0204802), the National Natural Science Foundation of China (31930053, T2421004).

## Author Contributions

X.-Y.Y. designed the experiments, collected and analyzed the data in all experiments, and wrote the manuscript.

C.Z. collected and analyzed the data in Experiment 5.

Z.Z. collected and analyzed the data in Experiment 5.

A.L. and H.L. reviewed the manuscript and provided critical feedback.

H.Q. collected the data in Experiments 1 and 2.

F.F. supervised the study, designed the experiments, and wrote and reviewed the manuscript.

## Data and code availability

Data and code are available upon request.

