## Supplementary Information for "Tetanizing wakeful consolidation: ten-hertz repetitive visual stimulation enhances the offline gain of visual learning"

#### Supplementary methods

##### Participant screening criteria

All participants included had met the following screening criteria: 1) no participation in any other study during participation in the current study, 2) having normal or corrected-to-normal vision, 3) having no history or family history of any psychological or neurological disease, 4) having no metal implant, no claustrophobia or attention deficit and hyperactivity disorder (ADHD), 5) having regular sleeping routine and no habit of siesta, and 6) not during the period of ovulation or pregnancy. Questionnaires (Mandarin edition) used in screening included 1) the Epworth Sleepiness Scale[1], 2) the Depression Anxiety Stress Scales-21[2], 3) the Positive and Negative Affect Schedule[3], 4) the Beck Depression Inventory-II[4], 5) the Pittsburgh Sleep Quality Index[5], and 6) the Morningness Eveningness Questionnaire[6].

##### Criteria of eye movement

Epochs that met either one of the following criteria were considered contaminated by eye movement and were excluded from further analyses: 1) having an extremum VEOG value exceeding 20 μv from zero, 2) having a detected saccade or a period of unobserved pupil in the eye tracker data.

##### Luminance and contrast control

Both monitors used in the study was calibrated to a linear 0-100 cd·m$^{-2}$ luminance. Therefore, the contrast of visual stimuli was calculated as:

$$contrast = \frac{max - 50}{100 - 50} = \frac{50 - min}{50 - 0}$$

Here, *max* is the maximum luminance (cd·m$^{-2}$) of the stimulus (either a gabor or a grating), and *min* is the minimum stimulus luminance (cd·m$^{-2}$). The medium luminance of stimuli was fixed at 50 cd·m$^{-2}$.

**Fitting learning rate in Experiments 1 and 2**

In order to capture the learning rate of individual participants, we fitted the block-to-block discrimination thresholds to a power function:

$$\theta_i = \lambda \times i^{-\rho} + \beta$$

Here, $\theta_i$ denotes the discrimination threshold of the $i^{th}$ block, $\lambda$ is the multiplication factor, and $\beta$ is the intercept factor. The learning rate is denoted by $\rho$. Positive values of $\rho$ denote improvement while negatives denote impairment. A larger positive $\rho$ means faster learning.

### Supplementary Results

**Ten-Hz RVS did not augment the N1 component in the ipsilateral hemisphere**

As we previously showed, the 10-Hz RVS increased the negativity of the N1 component in the contralateral hemisphere. Here we demonstrated that the RVS yielded no effect on the N1 component in the ipsilateral hemisphere using a repeated measure ANOVA. We analyzed the peak-to-peak amplitude of N1 from the ipsilateral electrode (either P5 or P6, depending on the condition that the participant was under). We did not find significant main effects (session: ($F_{(1,42)} = 3.341$, $p = 0.075$, partial $\eta^2 = 0.074$); group: ($F_{(2,42)} = 0.899$, $p = 0.415$, partial $\eta^2 = 0.041$)). Neither did we find an interaction ($F_{(2,42)} = 0.082$, $p = 0.922$, partial $\eta^2 = 0.004$). Analyses of simple main effects showed that the ipsilateral N1 amplitude was not altered by the

RVS (0-Hz: ($F_{(1,42)} = 0.957$, $p = 0.334$, partial $\eta^2 = 0.022$); 10-Hz: ($F_{(1,42)} = 1.881$, $p = 0.178$, partial $\eta^2 = 0.043$; 1-Hz: ($F_{(1,42)} = 0.666$, $p = 0.419$, partial $\eta^2 = 0.016$. Fig. S1).

**Ten-Hz RVS resulted in transient increase in alpha power in the occipital lobe**

Besides ERP analysis, we investigated flicker-induced neural oscillatory activities in the 10-Hz RVS group. Preprocessing was similar to that in “EEG” of “Methods”. Here, epochs were extracted from 1 second before to 2 seconds after the onset of the first grating of the 10-grating sequence (10 gratings constituted one flicker and extended a period of 1 second). The onset of a flicker was defined as the onset of the first grating, and the offset of a flicker was defined as the offset of the last ($10^{th}$) grating. Therefore, epochs extended from 1 second before flicker onset to 1 second after flicker offset. Data for one participant was excluded due to trigger malfunctions. Epochs from electrodes O1, PO3, and PO7 (or O2, PO4, and PO8) were averaged for each participant and then fitted to a sinusoidal function with a frequency fixed at 10 Hz, as was the RVS frequency in this group. Results revealed that the 10-Hz RVS transiently increased alpha activity power during the flickers and that the increase was more potent in the contralateral occipital sites compared to the ipsilateral sites (paired t-test: $t_{(13)} = 4.37$, $p < 0.001$; Fig. S2).

In Experiment 4, participants were instructed to rest with their eyes closed for 50 seconds between every two consecutive RVS blocks. We recorded EEG signals during this break and analyzed one period immediately before the RVS was implemented (Epoch 0) and one period after each RVS block (Epochs 1 to 5), resulting in a total of 6 periods. Epochs were extracted as from the $5^{th}$ to the $40^{th}$ second of each 50-second period and were baseline-corrected. Epochs with significant eye movements were excluded from further analyses. We performed fast Fourier transformation and spectrum normalization. Results showed that alpha oscillation was most significant in the occipital lobe (Fig. S3). We then selected the electrodes O1, PO3, PO7 (or O2,

PO4, PO8) and calculated their mean spectrum as oscillation activity in either the contralateral or ipsilateral occipital area. Alpha power was defined as the mean power of 3 different frequencies (9, 10, and 11 Hz). We then analyzed the effect of the RVS on alpha power using linear mixed effect models (group × epoch). We did not observe a significant effect of group ($F_{(2,74)} = 1.217$, $p = 0.302$) or epoch ($F_{(1,189)} = 2.647$, $p = 0.105$), or a significant interaction ($F_{(2,189)} = 1.770$, $p = 0.173$). These results demonstrated that, though significant alpha activity was detected in the occipital lobe, the 10-Hz RVS did not entrain alpha oscillation in the occipital lobe that lasted till after the offset of the 10-Hz RVS (Fig. S4).

### Supplementary figures

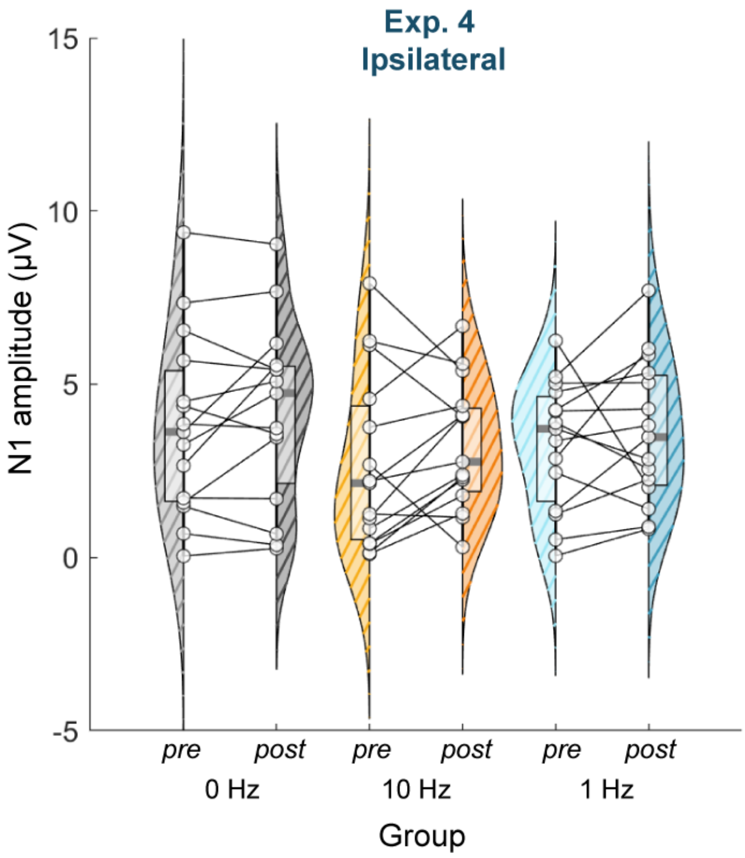


**Fig. S1. Amplitudes of N1 component extracted from the ipsilateral electrode.** The open circles denote individual participants. Filled curved areas, transparent squares, and the horizontal lines within the squares indicate scaled probability density estimates, the 25-75% quantiles, and medians, respectively.

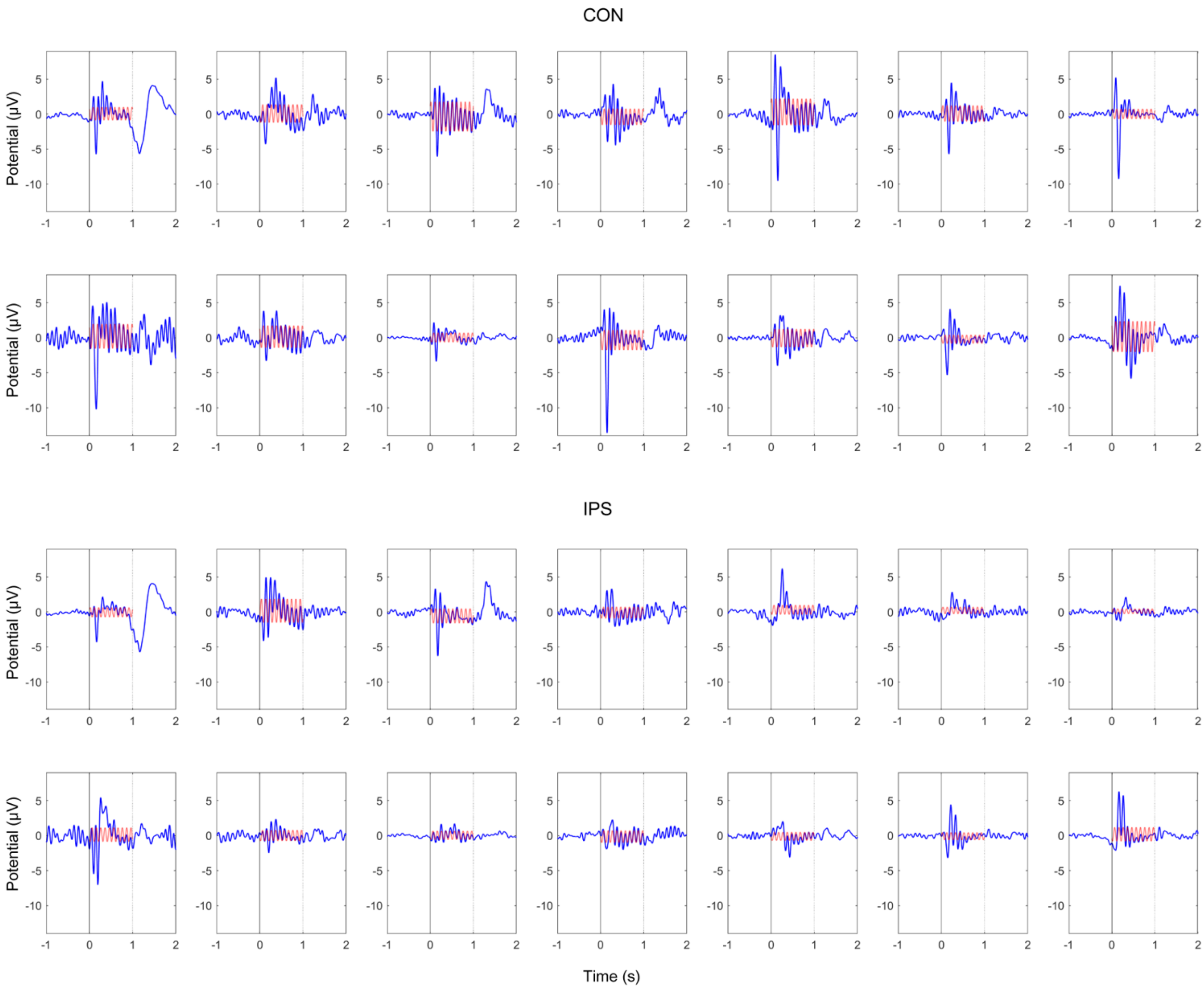


**Fig. S2. Transient increase in alpha power by the 10-Hz RVS in the occipital lobe.** Individual averaged-across-trials potentials (N = 14) are depicted. The upper and lower panels show data from occipital electrodes contralateral and ipsilateral to the RVS stimuli, respectively. Data and functions are separately plotted in blue and red. Vertical lines represent 10-Hz flickers' onset (solid) and offset (dotted).

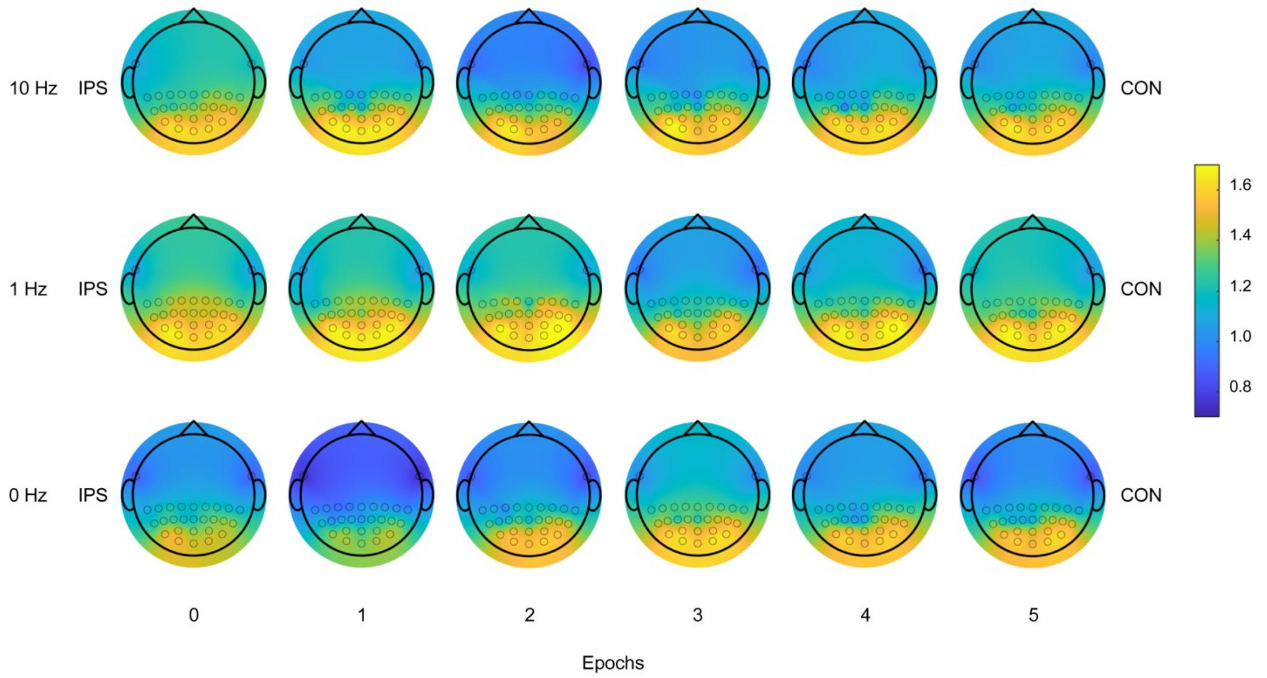


**Fig. S3. Fieldmaps of alpha power distribution.** EEG signals were selected from 26 posterior electrodes and were segmented into 35-second epochs within between-block rest periods. Alpha oscillations were significant in the occipital lobe across all groups and epochs.

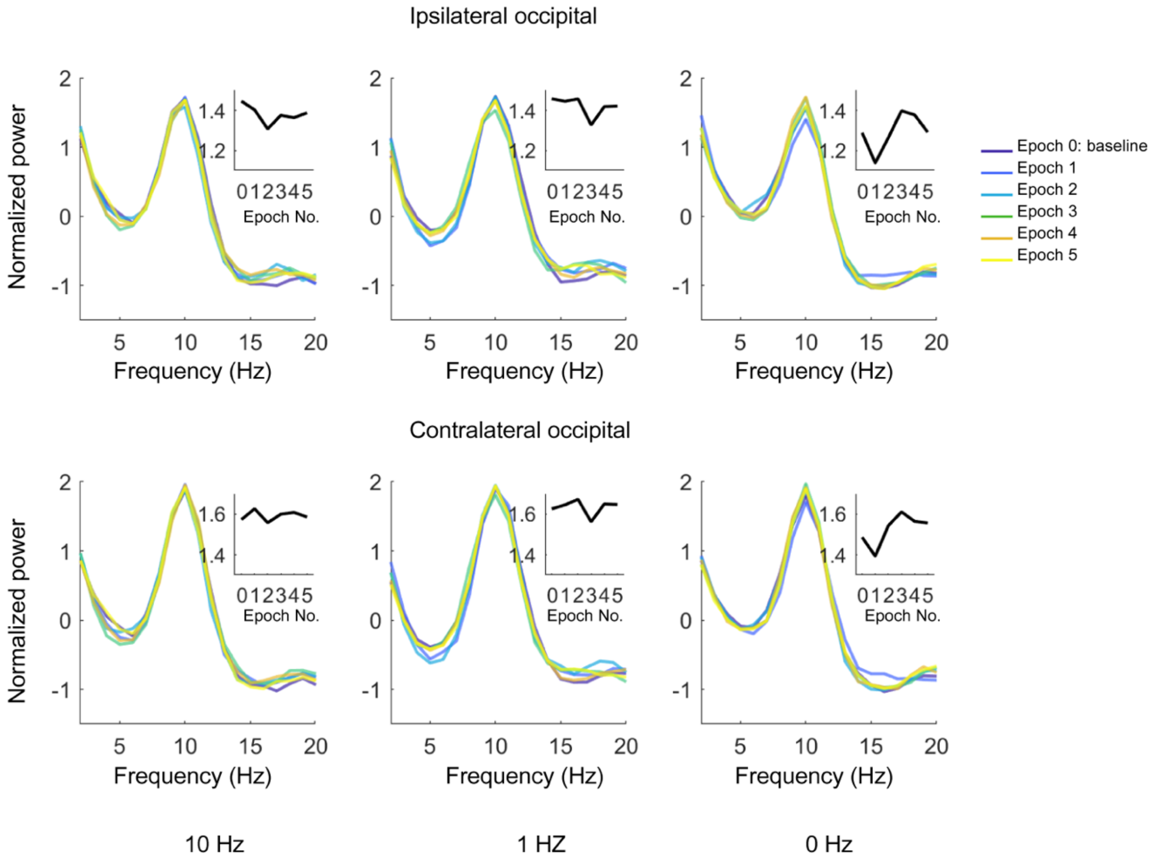


**Fig. S4. Spectra of occipital neural oscillatory activities.** The normalized spectra of each group are plotted. Inserted are the changes in alpha power. Upper and lower panels are spectra calculated from the ipsilateral and contralateral occipital electrodes, respectively. Notably, alpha power in the contralateral occipital lobe was stronger than that in the ipsilateral lobe.
